# The CD73 immune checkpoint promotes tumor cell metabolic fitness

**DOI:** 10.1101/2022.11.29.518350

**Authors:** David Allard, Isabelle Cousineau, Eric Ma, Bertrand Allard, Yacine Barèche, Hubert Fleury, John Stagg

## Abstract

CD73 is an ectonucleotidase overexpressed on tumor cells that suppresses anti-tumor immunity. Accordingly, several CD73 inhibitors are currently being evaluated in the clinic, including in large randomized clinical trials. Yet, the tumor cell-intrinsic impact of CD73 remain largely uncharacterized. Using metabolomics, we discovered that CD73 significantly enhances tumor cell mitochondrial respiration and aspartate biosynthesis. Importantly, rescuing aspartate biosynthesis was sufficient to restore proliferation of CD73-deficient tumors in immune deficient mice. Seahorse analysis of a large panel of mouse and human tumor cells demonstrated that CD73 enhanced oxidative phosphorylation (OXPHOS) and glycolytic reserve. Targeting CD73 decreased tumor cell metabolic fitness, increased genomic instability and suppressed poly ADP ribose polymerase (PARP) activity. Our study thus uncovered an important immune-independent function for CD73 in promoting tumor cell metabolism, and provides the rationale for previously unforeseen combination therapies incorporating CD73 inhibition.

## Introduction

CD73 is an ectonucleotidase that catalyzes the phosphohydrolysis of adenosine monophosphate (AMP), thus contributing to the breakdown of pro-inflammatory adenosine triphosphate (ATP) into immunosuppressive ADO. Accumulation of extracellular ADO in the tumor microenvironment favors cancer progression by activating ADO specific G-coupled protein receptors (GPCRs) expressed on immune cells, endothelial cells, fibroblasts and tumor cells. Of the four ADO receptors, high affinity A2A and low affinity A2B receptors elevate intracellular cyclic AMP (cAMP) levels and contribute to suppress immune cell function. In human cancers, such as triple negative breast cancer (TNBC) and ovarian cancer, CD73 expression is generally associated with poor clinical outcomes, impaired tumor immune surveillance and resistance to chemotherapy (1, 2). Potent and selective CD73 inhibitors have recently been developed and are now being evaluated in randomized clinical trials with encouraging preliminary results reported in lung cancer and pancreatic cancer patients (3, 4).

While CD73 inhibitors can stimulate anti-tumor immunity, early studies reported that targeting CD73 can also inhibit human tumor xenografts in severely immunocompromised mice (5–8). Moreover, pharmacological inhibition or gene-deletion of CD73 has also been observed to increase the sensitivity of CD73-expressing tumor cells to cytotoxic drugs in vitro (1, 9, 10). Altough activation of pro-survival pathways downstream of A2B receptors has been proposed to explain immune-independent effects of CD73, alternative hypotheses have not been explored (11, 12).

Rewiring of cancer cells’ metabolism is central to their adaptation to hostile tumor microenvironments (7). While cancer cells can shift their metabolism from highly efficient ATP-producing oxidative phosphorylation (OXPHOS) to low ATP-yielding glycolysis despite presence of oxygen (e.g., Warburg effect), mitochondrial respiration has been shown to be critical for proliferating cells and for maintaining tumor cells invasiveness, metabolic adaptation, chemoresistance and stemness. Accordingly, tumor cells are not only capable of respiration but often require respiration for survival (13).

In proliferating cells, mitochondrial respiration provides electron acceptors to support biosynthesis of aspartate, an indispensable amino acid for nucleotide and protein synthesis. Mitochondrial respiration requires nicotinamide adenine dinucleotide (NAD) in its oxidized and reduced forms to allow movement of electrons through the electron transport chain (ETC). In addition of supporting respiration, NAD is also an important co-substrate for many enzymes, notably poly-ADP-ribose polymerases (PARPs) that play a central role in genomic stability by catalyzing the addition of poly-ADP-ribose to proteins (also known as PARylation) (14, 15).

Intracellular NAD levels are maintained either by *de novo* synthesis from L-tryptophan (Trp), or by more effective salvage pathways that recycle NAD from nicotinamide (NAM), nicotinamide mononucleotide (NMN) or nicotinamide riboside (NR) (16). Extracellular NMN is dephosphorylated to NR before cellular internalization by equilibrative nucleotide transporters (ENTs). Intracellular NR is then re-phosphorylated by NRK1 into NMN, a substrate for adenylyl transferases (NMNATs) to generate NAD (17, 18). NMN can also be generated from intracellular or extracellular NAM via the rate-limiting enzyme NAMPT (18). Interestingly, some studies have suggested a role for CD73 in NAD synthesis from extracellular NR (19, 20). Yet, the impact of CD73 on tumor cell metabolic pathways remains unknown (21–22).

We here investigated metabolic consequences of targeting CD73 in a panel of human and murine cancer cells. We observed that CD73 deficiency or pharmacological inhibition induces an important aspartate-dependent metabolic vulnerability in tumor cells characterized by suppressed mitochondrial respiration and increased genomic instability. Our study thus provides a rationale for exploring novel combination strategies with anti-CD73 therapies, such as PARP inhibition and mitochondria targeting agents.

## Results

### CD73 favors tumor growth independently of its immunosuppressive function

We evaluated the impact of tumor cell-associated CD73 by transiently transfecting a CRISPR-Cas9 construct into human and mouse tumor cell lines followed by flow cytometry-based sorting of polyclonal CD73-negative (CD73neg) and CD73-positive (CD73pos) fractions (**Fig. S1A-E**). Consistent with prior work (8) and in support of immune-independent effects, CD73 deletion in MDA-MB-231 human triple negative breast cancer (TNBC) cells significantly suppressed tumor growth in severely immunodeficient Nod-Rag-gamma (NRG) mice, which lack T cells, B cells, NK cells and functional macrophages (**Fig. 1A**). To further characterize the immune-independent effects of CD73, we next compared the in vitro proliferation rates of CD73pos and CD73neg MDA-MB-231 cells. While CD73 gene deletion had no impact in standard cell culture conditions (i.e. 25 mM glucose and 2 mM glutamine) (**Fig. 1B, Fig. S2)**, it significantly suppressed cell proliferation in low glucose (0-2.5 mM) and low glutamine (0-0.1 mM) conditions (**Fig. 1B, Fig. S2**), despite no difference in cellular uptake of glucose or glutamine, as shown using a fluorescent glucose analog (i.e. 2-NDBG; **Fig. 1C**) and isotope-labelled glutamine (i.e. U-[^13^C]-glutamine; **Fig. 1D**).

**Figure 1.**
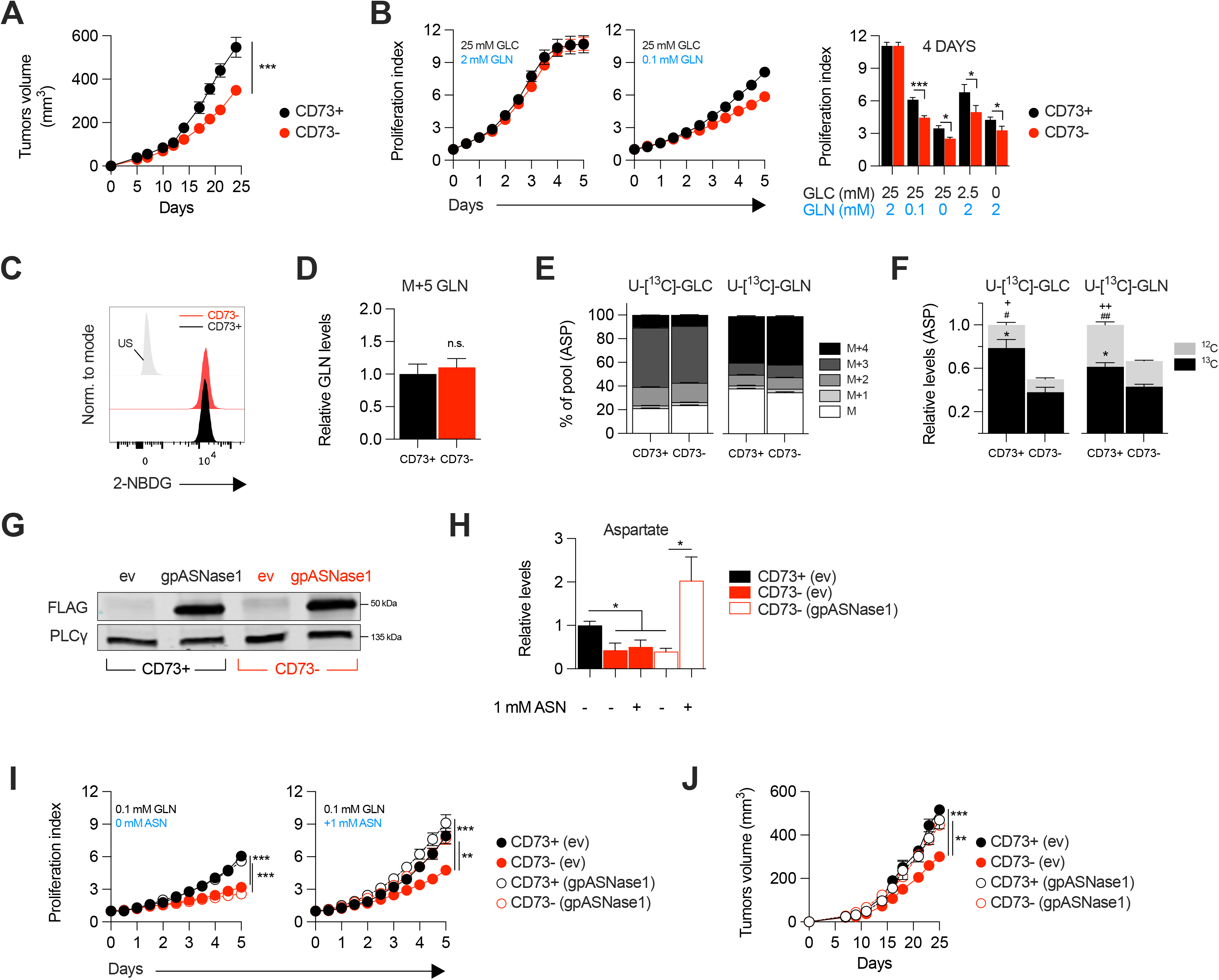
CD73 promotes tumor growth independently from immune suppression through aspartate biosynthesis. **A** Sub-cutaneous tumor growth of CD73pos (CD73+) and CD73neg (CD73-) MDA-MB-231 tumors in immune-deficient NRG mice. Results shows combined data of 2 independent experiments cumulating 15 mice per groups. **B** In vitro proliferation of CD73+ and CD73-MDA-MB-231 cells was analyzed by Incucyte live imaging technology in presence of excess or restricted conditions of glucose (GLC) and glutamine (GLN). Data from representative proliferation curves (left panels) and pooled quantification of proliferation after 4 days of culture (right panel) are shown (n=3). **C** FACS analysis of in vitro incorporation of a fluorescent glucose analog (2-NBDG) in CD73+ and CD73-MDA-MB-231 cells (n=3). US=unstained control. **D** GC-MS analysis of in vitro incorporation of a stable isotopically labeled glutamine tracer in vitro (U-[^13^C]-GLN). M+5 glutamine (GLN) levels were normalized on cell numbers and are shown relative to CD73+ cells (n=1). **E** Incorporation of isotopically labeled glucose (U-[^13^C]-GLC; left panel) or glutamine (U-[^13^C]-GLN; right panel) into aspartate (ASP) compared between CD73+ and CD73-MDA-MB-231 cells (n=1). **F** SITA analysis of glucose (left panel) and glutamine (right panel) contribution to ASP intracellular biosynthesis (n=1). ASP levels were normalized on cell number and are shown relative to CD73+ cells. Stats: ^*^comparison of ^13^C levels; ^#^Comparison of ^12^C levels; ^+^Comparison of total ^13^C+^12^C levels. **G** Western blot of CD73+ and CD73-MDA-MB-231 cells expressing empty vector (ev) or FLAG-tagged gpASNase1. PLCγ is shown as a loading control. **H** Intracellular ASP levels measured in cells expressing or not (ev) the gpASNase1 upon exposure to exogenous ASN (1mM) for 24h. ASP levels were normalized on cell number and are shown relative to CD73+ (ev) cells (n=1). **I** In vitro proliferation of gpASNase1-expressing and non-expressing (ev) MDA-MB-231 cells in presence (right panel) or not (left panel) of exogenous ASN (n=2). **J** In vivo tumor growth of 2×10^6^ gpASNase1-expressing or not (ev) CD73+ and CD73-MDA-MB-231 cells in NRG mice (CD73+ (ev) n=5; CD73- (ev) n=7; CD73+ (gpASNase1) n=8; CD73- (gpASNase1) n=8). Means +/- SEM are shown (*p<0.05; **p<0.01; ***p<0.001 by Student T tests).

### CD73 enhances aspartate biosynthesis to promote tumor growth

We hypothesized that CD73 was involved in promoting tumor cell metabolism. To test this, we performed stable isotope tracer analysis (SITA) of labeled glucose or glutamine in CD73pos and CD73neg MDA-MB-231 cells. Strikingly, while the distribution and relative abundance of most intermediate metabolites were unaltered (**Fig. S3A-D**), aspartate biosynthesis from glucose or glutamine (**Fig. 1E**) was significantly reduced – by nearly 50% – in CD73neg tumor cells (**Fig. 1F**).

To assess whether this decrease in aspartate biosynthesis was responsible for the suppressed growth of CD73-deficient tumor cells, we tested the ability of aspartate supplementation to restore proliferation of CD73neg MDA-MB-231 cells. We used a strategy developed by Sullivan et al. (23), whereby an asparaginase enzyme (i.e. gpASNase1) is introduced into cells to restore intracellular aspartate levels upon exposure to exogenous asparagine (since exogenous aspartate cannot be readily incorporated into cells). Stable expression of gpASNase1 (**Fig. 1G**) and asparagine treatment indeed significantly increased intracellular aspartate levels (**Fig. 1H**) and effectively restored proliferation of CD73neg MDA-MB-231 tumor cells, both *in vitro* in glutamine-restricted conditions (**Fig. 1I**) and *in vivo* in immunodeficient NRG mice (**Fig. 1J**). Taken together, our results demonstrated that CD73 promoted tumor growth by increasing aspartate biosynthesis.

### CD73 regulates metabolic fitness of cancer cells

Since aspartate biosynthesis is an essential function of mitochondrial respiration (24–26), and that aspartate fuels OXPHOS (27), we evaluated the impact of CD73 on mitochondrial respiration. For this, we measured oxygen consumption rate (a surrogate of OXPHOS) and extracellular acidification rate (a surrogate of glycolysis) in response to standard mitochondrial stress tests (**Fig. 2A**). Strikingly, CD73neg MDA-MB-231 cells displayed significantly reduced OXPHOS (**Fig. 2B**) and reduced glycolytic reserve compared to CD73pos cells (**Fig. 2C**). In addition, blocking CD73 with a neutralizing mAb (28) also significantly suppressed OXPHOS and glycolytic reserve (**Fig. 2B-C**). Importantly, similar results were obtained in additional murine cell lines (4T1, SM1 LWT1), human cell lines (SK23MEL, PANC1, SKOV3 and HCC1954) and in primary murine embryonic fibroblasts (MEF) derived from CD73-deficient mice (**Fig. 2D-H, Fig. S1F**).

**Figure 2.**
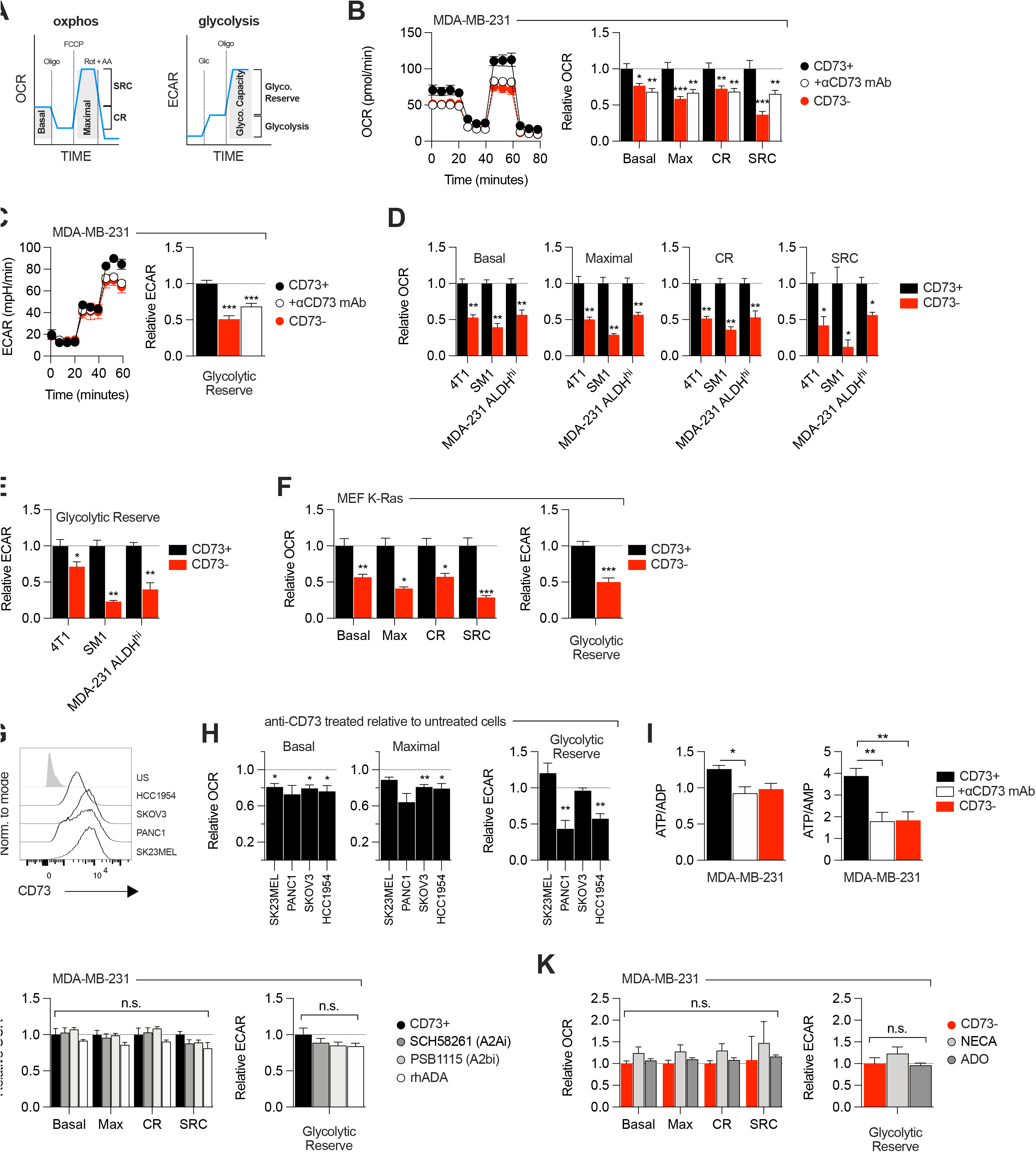
CD73 regulates metabolic fitness of cancer cells independently from adenosine signaling. **A** Schematized metabolic profiling and parameters generated on a Seahorse analyzer. OXPHOS and glycolytic parameters were calculated using Agilent calculator tool. **B** OXPHOS profile (left panel) and parameters (right panel) of CD73pos (CD73+; +/- 1μg/mL anti-CD73 mAb 7G2 for 48h) and CD73neg (CD73-) MDA-MB-231 cells. OXPHOS parameters are shown relative to CD73+ cells from pooled experiments (n=3). **C** Glycolytic profile (left panel) and glycolytic reserve (right panel) of CD73+ (+/- 1μg/mL anti-CD73 mAb 7G2 for 48h) and CD73 - MDA-MB-231 cells. Glycolytic reserve is shown relative to CD73+ cells from pooled experiments (n=3). **D-E** Oxphos parameters (**D**) and glycolytic reserve (**E**) compared between various CD73-expressing and CD73 CRISPR-knockout cell lines. ALDHhi cells were sorted from MDA-MB-231 cells using the Aldefluor kit. Data are shown relative to CD73+ cells (MDA-231 ALDHhi n=1, 4T1 and SM1 LWT1 n=2). **F** OXPHOS parameters (left panel) and glycolytic reserve (right panel) compared between primary MEF isolated from WT and CD73^-/-^ C57BL/6 mice and transformed with an oncogenic K-Ras. Data are shown relative to WT cells (n=2). **G** CD73 expression profile on various human cancer cell lines analyzed by FACS. US=unstained control. **H** OXPHOS parameters (left and middle panels) and glycolytic reserve (right panel) of anti-CD73 mAb (7G2)-treated cells compared to untreated cells. Data are shown relative to respective untreated cells (n=1 per cell line). **I** LC-MS analysis of total intracellular levels of ATP/ADP and ATP/AMP ratios in CD73+ (+/- 1μg/mL anti-CD73 mAb 7G2 for 48h) and CD73-MDA-MB-231 cells (n=2). **J** OXPHOS parameters (left panel) and glycolytic reserve (right panel) of CD73+ MDA-MB-231 cells treated with either A2A inhibitor SCH 58261, A2B inhibitor PSB 1115 or human recombinant ADA. Data are shown relative to untreated CD73+ cells (A2ai and A2bi n=2, rhADA n=1). **K** OXPHOS parameters (left panel) and glycolytic reserve (right panel) of NECA- or ADO-treated CD73-MDA-MB-231 cells. Data are shown relative to untreated CD73-cells (NECA n=2, ADO n=1). Means +/- SEM are shown (*p<0.05; **p<0.01; ***p<0.001 by Student T tests).

Consistent with the observed defect in mitochondrial respiration, CD73 gene-targeted tumor cells also displayed reduced ratios of ATP to ADP or AMP (**Fig. 2I**). However, no difference was observed in GOT1 and GOT2 mRNA expression, which encode enzymes that generate aspartate from oxaloacetate (24) (**Fig. S3E**), or other tricarboxylic acid (TCA) cycle-related enzymes (**Fig. S3E**).

As CD73 is overexpressed in tumor-initiating cells (TIC) to promote stemness (29, 30), and that TIC rely on OXPHOS to support tumorigenesis (31), we next evaluated the impact of CD73 on the metabolic fitness of putative TIC (ALDH^High^ CD44^+^ CD24^-^) derived from MDA-MB-231 cell cultures (**Fig. S1G**). Similar to the bulk culture, CD73neg TIC also displayed significantly suppressed OXPHOS and reduced glycolytic reserve compared to CD73pos TIC (**Fig. 2D-E**). Taken together, our data demonstrated that CD73 plays a critical role in promoting mitochondrial respiration in proliferating tumor cells.

### ADO-independent metabolic functions of CD73

We next evaluated whether the metabolic functions of CD73 were mediated by ADO receptor signaling. MDA-MB-231 cells express A2A and A2B receptors (32). Nevertheless, treatment with an A2A antagonist (SCH58261) or an A2B antagonist (PSB1115) had no effect on mitochondrial respiration or glycolysis (**Fig. 2J**). To rule-out a potential role for other ADO receptors, we treated MDA-MB-231 cells with recombinant adenosine deaminase (rhADA) or the pan-ADO receptor agonist NECA. Depletion of extracellular ADO with rhADA (**Fig. 2J**), or treatment with NECA (**Fig. 2K**), also had no impact on mitochondrial respiration or glycolysis of MDA-MB-231 cells. Our results thus indicated that CD73 promoted metabolic fitness independently of ADO signaling.

### CD73 contributes to metabolic fitness of cancer cells through NAD synthesis

NAD+ is a coenzyme that plays a central role in mitochondrial respiration (16). We thus investigated whether CD73 could regulate intracellular NAD levels. Using mass-spectrometry (LC-MS), we observed that CD73 gene-targeting or antibody-mediated enzymatic inhibition was associated with a significant reduction of intracellular NAD+ levels in MDA-MB-231 (**Fig. 3A**) or 4T1.2 cells (**Fig. S4A**). Depleting extracellular ADO with exogenous rhADA had no effect on intracellular NAD+ levels, further supporting an ADO-independent mechanism (**Fig. 3A**).

**Figure 3.**
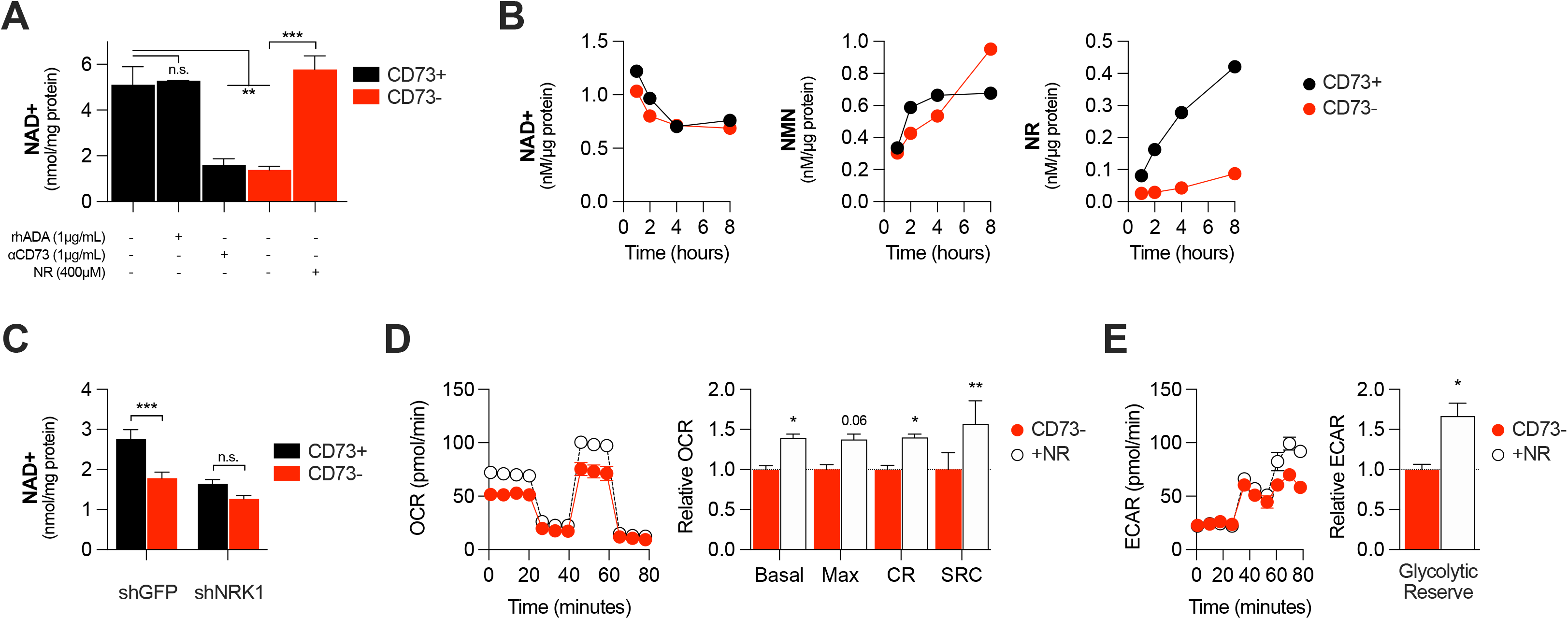
CD73 contributes to metabolic fitness of cancer cells through NAD synthesis. **A** LC-MS analysis of intracellular nicotinamide adenine levels (NAD+) normalized to protein content of CD73pos (CD73+; +/- 1μg/mL αCD73 mAb or 1μg/mL rhADA for 48h) and CD73neg (CD73-; +/- 100μM NR supplemented 2x per day for 48h) MDA-MB-231 cells (n=2). **B** LC-MS analysis of endogenous extracellular production of NAD+ (left panel), NMN (middle panel) and NR (right panel) in supernatant of CD73+ and CD73-MDA-MB-231 cells. Levels were normalized on protein content of adherent cells (n=1). **C** LC-MS analysis of intracellular nicotinamide adenine levels (NAD+) normalized to protein content of CD73+ and CD73-MDA-MB-231 cells transfected with either shGFP or shNRK1 (n=2). **D** Oxygen consumption rate (OCR) profile (left panel) and OXPHOS parameters (right panel) of CD73-MDA-MB-231 cells cultured with 400μM NR for 48h (n=3). **E** Extracellular acidification rate (ECAR) profile (left panel) and glycolytic reserve (right panel) of CD73 - MDA-MB-231 cells cultured with 400μM NR for 48h (n=3). Means +/- SEM are shown (*p<0.05; **p<0.01; ***p<0.001 by 1-way ANOVA (**A, C**) or Student T tests (**D-E**)).

We next tested whether CD73 can serve tumor cells to generate NR, a precursor for NAD synthesis. Using purified recombinant human CD73 and mass-spectrometry, we confirmed hydrolysis of exogenous NMN to NR by CD73 (**Fig. S4B**). We then tested the impact of CD73 on the extracellular accumulation of NAD+, NMN and NR from MDA-MB-231 cell cultures. Strikingly, we observed that MDA-MB-231 cell cultures accumulated extracellular NMN and rapidly hydrolyzed it to NR in a CD73-dependent manner (**Fig. 3B**). In contrast, CD73 had no impact on extracellular NAD+ levels (**Fig. 3B**).

NR must be phosphorylated by intracellular NRK1 to promote intracellular NAD synthesis. We thus tested whether NRK1 was involved in CD73-mediated NAD synthesis. In support of a metabolic role for CD73-mediated hydrolysis of extracellular NMN to NR, shRNA-mediated knockdown of NRK1 (**Fig. S4C**) prevented CD73 from increasing intracellular NAD+ levels in MDA-MB-231 cells (**Fig. 3C**).

To further assess the importance of NR production for CD73 function, we tested whether exogenous NR could rescue the metabolic dysfunctions of CD73neg tumor cells. Indeed, treatment of CD73neg MDA-MB-231 tumor cells with exogenous NR for 48 hours significantly increased their oxygen consumption (**Fig. 3D**) and glycolytic reserves (**Fig. 3E**). Altogether, our results support a model whereby CD73-dependent extracellular NR promotes intracellular NAD biosynthesis, which in turn enhances mitochondrial respiration and aspartate biosynthesis.

### CD73 deficiency decreases PARP activity and sensitizes cancer cells to DNA-damaging agents

In addition of being essential for mitochondrial respiration, NAD is also an important co-factor for PARP enzymes that maintain genomic stability by promoting DNA repair. Interestingly, early studies reported that tumor-derived CD73 can promote chemoresistance (1, 9, 10), although the exact mechanism has remained elusive. We hypothesized that by increasing intracellular NAD levels, CD73 may regulate PARP activity and therefore promote genomic stability. To assess this, we measured PARylation levels by western blotting in H2O2-treated MDA-MB-231 cells. As shown in Fig. 4, CD73-deficient tumor cells displayed significantly reduced PARylation compared to CD73-proficient tumor cells (**Fig. 4A**). Moreover, targeting CD73 with a neutralizing mAb also reduced PARylation levels, and NR supplementation rescued PARylation in CD73neg cells (**Fig. 4B-C**). Similar results were obtained in UWB1.289 human ovarian cancer cells (**Fig. S5A-B**).

**Figure 4.**
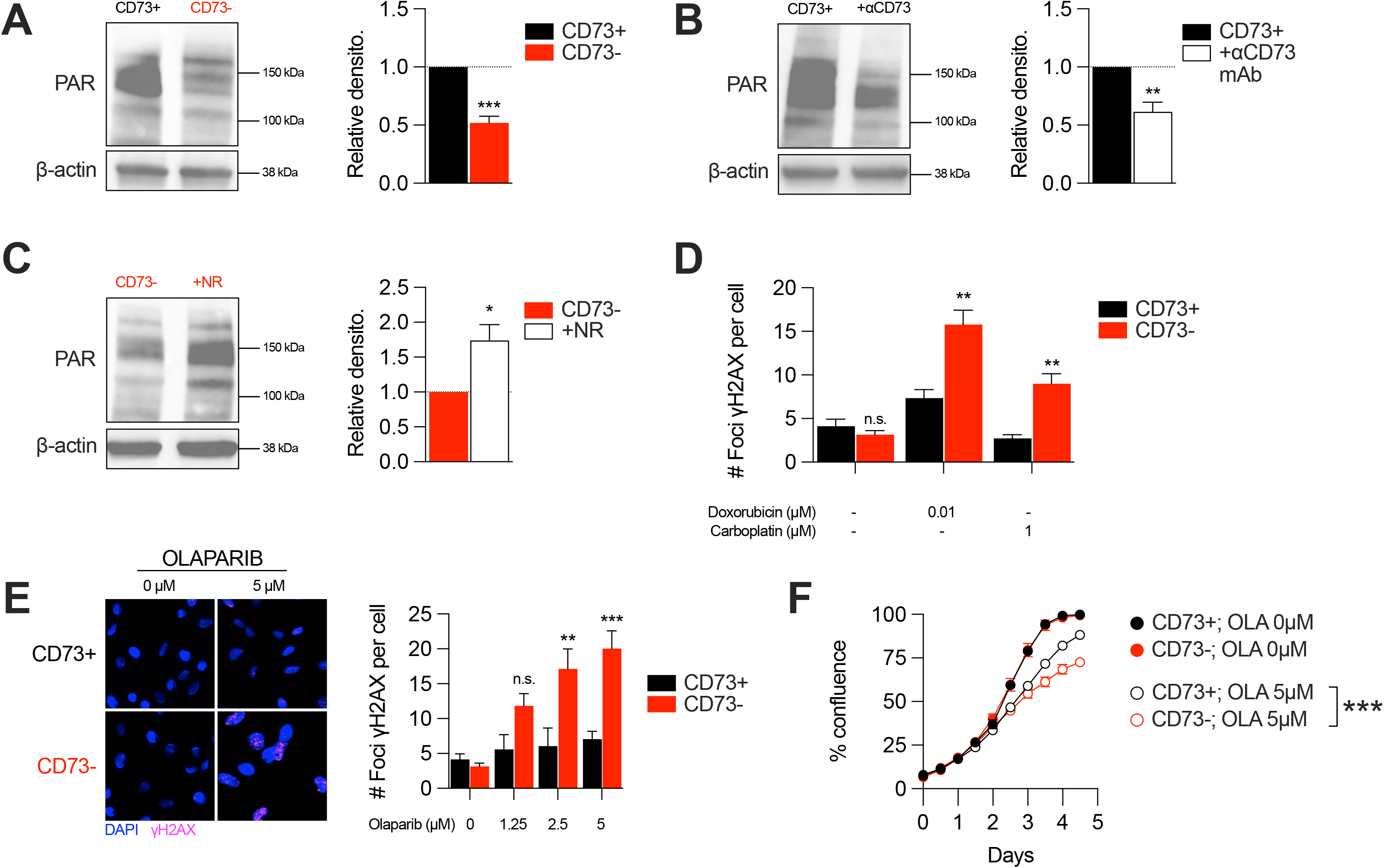
CD73 deficiency decreases PARP activity and sensitizes cancer cells to DNA-damaging agents. **A** Western blot (left panel) showing PARylation levels in CD73pos (CD73+) and CD73neg (CD73-) MDA-MB-231 cells. Densitometry (right panel) is shown relative to CD73+ cells (n=5). **B** Western blot (left panel) showing PARylation levels in CD73+ MDA-MB-231 cells treated with αCD73 mAb (1μg/mL; clone 7G2; 48h). Densitometry (right panel) is shown relative to untreated CD73+ cells (n=2). **C** Western blot (left panel) showing PARylation levels in CD73-MDA-MB-231 cells treated with NR (400μM; 48h). Densitometry (right panel) is shown relative to untreated CD73-cells (n=2). **D** Number of γH2AX foci per CD73+ and CD73-MDA-MB-231 cells treated with doxorubicin (0.01 μM) or carboplatin (1μM) for 48h (n=2). **E** γH2AX staining of CD73+ and CD73-MDA-MB-231 cells treated with olaparib (0-5μM) for 48h. Left panels show representative images for γH2AX (magenta) and DAPI (blue) staining. Data show number of γH2AX foci per cells (right panel; n=2). **F** In vitro proliferation of CD73+ and CD73-MDA-MB-231 cells treated with (0-5μM) olaparib for 4 days. Confluence was measured using an Incucyte instrument (n=1). Means +/- SEM are shown (*p<0.05; **p<0.01; ***p<0.001 by Student T tests).

Given the impact of CD73 on PARylation, we next compared the levels of DNA damage in CD73pos and CD73neg tumor cells in response to cytotoxic drugs for TNBC or ovarian cancer. Strikingly, treatment of CD73-deficient MDA-MB-231 or UWB1.289 cells with doxorubicin and carboplatin was associated with significantly greater DNA damage compared to CD73-proficient cells (**Fig. 4D & S5C**). Since decreasing intracellular NAD levels has also been shown to sensitize tumor cells to PARP inhibitors (33), we also evaluated the impact of CD73 on olaparib activity. Accordingly, CD73-deficient MDA-MB-231 cells were significantly more sensitive to olaparib activity in vitro (**Fig. 4E-F**). Altogether, our results suggest that by regulating intracellular NAD levels, CD73 plays an important role in promoting PARP activity and maintaining genomic stability of tumor cells.

## Discussion

In this study, we demonstrated that the ectonucleotidase CD73, often overexpressed on tumor cells and considered as an important cancer immune checkpoint, enhances tumor cell metabolic fitness in a cell-autonomous manner. CD73 promotes mitochondrial respiration, which generates aspartate required for nucleotide and protein synthesis, and maintains genomic stability notably by increasing the activity of NAD+-dependent PARP enzymes. Surprisingly, these metabolic functions of CD73 were independent of ADO signaling, instead relying on the hydrolysis of extracellular NMN to NR and its metabolism by NRK1.

The importance of CD73 for intracellular NAD synthesis has been controversial (21, 22, 34). Using a colorimetric assay, Wilk et al. (22) observed no significant effect of inhibiting or deleting CD73 on intracellular NAD concentrations. These discrepancies from our own data may stem from the different sensitivity of the assays (colorimetric versus LC-MS), the different nature of the inhibitors and the different cell lines studied. Notably, Wilk et al. studied MCF-7 cells that express >10-fold lower levels of CD73 compared to the cell lines we analyzed. Importantly, we validated the impact of CD73 on mitochondrial respiration in a large panel of human and mouse tumor cell lines, as well as in primary fibroblasts derived from CD73 gene-targeted mice. Moreover, the fact that NR supplementation restored intracellular NAD+ content in CD73-deficient tumor cells, and that CD73 failed to increase NAD+ levels in the absence of NRK1 (since NRK1 is necessary and rate-limiting for the generation of NAD+ from exogenous NR (17)), further support the notion that tumor cell-associated CD73 contributes to maintain intracellular NAD pools.

Intriguingly, we observed that NMN accumulates in the extracellular milieu of proliferating MDA-MB-231 tumor cells. The mechanism leading to such extracellular NMN accumulation remains unclear. We ruled out the culture media as a potential source of NMN. Extracellular NMN may come from NAMPT activity (intracellular or extracellular) on nicotinamide (NAM) that reaches the extracellular milieu as a result of cell death or stress (18). NAD release may be another source of extracellular NMN via active or passive release mechanisms followed by consumption by glycohydrolases to generate extracellular NAM. Finally, although MD-MB-231 and 4T1.2 cells do not express CD38, we cannot exclude the involvement of other ecto-NADases.

In the tumor microenvironment, CD38 expression on immune cells may also contribute to consume extracellular NAD+. Production of NAM from NAD-consuming enzymes and NAMPT may thus lead to CD73-mediated NR production. Considering that CD38 can cooperate with CD73 for extracellular ADO production (35), it would be of interest to assess the impact of host CD38 on CD73-mediated tumor cell metabolic fitness. Of note, while some studies suggested that CD73 may hydrolyze extracellular NAD+ (19, 36), we and others (22) did not observe this.

Consistent with the observed decrease in intracellular NAD pools, and the importance of NAD for glycolysis (37–41), we found that CD73-deficient tumor cells also displayed impaired maximal glycolytic capacity following oligomycin treatment, which inhibits ATP synthase and forces cells to redirect pyruvate to lactate conversion via glycolysis (38).

Intriguingly, we also demonstrated that tumor-derived CD73 promoted DNA repair and was associated with increased PARP activity. Accordingly, targeting CD73 sensitized tumor cells to chemotherapy and PARP inhibition. Besides PARPs, other DNA repairing enzymes also require NAD+ (42), such as SIRT1 and SIRT6, which play a critical role in positively regulating homologous recombination (HR) and nonhomologous end joining (NHEJ) pathways. It would therefore be of interest to further evaluate the impact of CD73 on sirtuin-mediated HR and NHEJ. A broad impact of CD73 on DNA repair mechanisms might explain the fact that CD73-deficient tumor cells become sensitive to PARP inhibition despite expressing BRCA. Accordingly, others have shown that targeting NAD+ metabolism (by blocking NAMPT) enhances the activity of olaparib in BRCA1-competent TNBC cells (33). Another possible explanation is that targeting NAD+ metabolism may decrease BRCA1 expression or activity, as was previously reported (43).

In conclusion, our study sheds new light on the immune-independent functions of CD73. We demonstrated that CD73 promote tumor growth in a cancer-cell autonomous manner by increasing metabolic fitness. This in turn favors cancer cell adaptation to nutrient-deprived microenvironments and cytotoxic stress by increasing OXPHOS, aspartate biosynthesis and DNA repair. Our study highlights new therapeutic opportunities that may be exploited in novel combination anti-cancer strategies.

## Acknowledgments

The authors thank Julien Lamontagne, Alexia Grangeon, Maxime Cahuzac, Dominique Gauchat, Daina Avizonis and Bozena Samborska for technical assistance.

## METHODS

Source and identifier of all reagents and resources are detailed in supplementary table 1.

### Cell lines generation and culture

CD73 gene-editing was generated by electroporation of all-in-one CRISPR/Cas9 vector (px330, Addgene) expressing the 20mer target sequence GCAGCACGTTGGGTTCGGCG (exon1), provided by Michael Hoelzel (University of Bonn, Germany). Cells were sorted based on CD73 expression (CD73+ and CD73-) within 1 week after transfection. Enzymatic functionality was verified using the commercially available malachite green kit assay (R&D system).

Empty vector (pLHCX-ev) and plasmid coding for a FLAG-tagged guinea pig asparaginase (pLHCX-gpASNase1) enzyme were kindly provided by Dr. Lucas B. Sullivan (Fred Hutchinson Cancer Research Center, Seattle, WA, USA) (23). CD73+ and CD73-MDA-MB-231 cells were infected with a retrovirus containing either pLHCX-ev or pLHCX-gpASNase1 and selected with 750μg/mL of Hygromycin B-containing media until all uninfected control cells died. For NRK1 knockdown in MDA-231 cells, CD73+ and CD73-cells were infected with a retrovirus containing either a control shRNA (shGFP) or shNRK1 and selected with 1μg/mL of puromycin-containing media until all uninfected control cells died. Plasmids were purchased from Sigma.

MEF were isolated from embryos at 14.5 days of gestation from WT and CD73^-/-^ C57BL/6 mice. Briefly, pregnant females were euthanized, uterus were dissected out and individual embryos separated from their yolk sacs into PBS-containing petri dishes. Limbs, tail, head and liver were cut off embryos before being digested and broken up by up-and-down pipetting in freshly thawed 0.25% trypsin solution and incubated at 37°C in water bath for 5 minutes cycles until only insoluble cartilage would remain and not settle down. Pooled dissociated cells were filtered through a 40μM mesh, centrifuged 10 minutes at 1200 rpm, resuspended in DMEM with 1% pen/strep antibiotics, seeded in 100 mm x 15 mm petri dishes (2 plates per embryo) and incubated in a 37°C/5% CO2 incubator. Growth media was replaced the next day and rapidly transformed within 7 days upon isolation with a lentiviral vector expressing an oncogenic K-Ras^G12V^ (pLenti PGK RasV12), kindly provided by Dr. Francis Rodier (Université de Montréal, Montréal, QC, CA) (44). Lentivirus was generated in HEK293FT cells by transfection using standard techniques and cells selected with 75μg/mL of hygromycin B until all uninfected cells were dead. Immortalized MEF clones upon K-Ras transformation were pooled and used for experimentation.

All cells were grown at 37°C in a 5% CO2 humid atmosphere. Cells were sub-cultured or media was changed every 3 days. MDA-MB-231, MDA-MB-231 ALDH^hi^, 4T1.2, SM1 LWT1, PANC-1 cell lines were cultured in antibiotics-free DMEM (Wisent) supplemented with 10% FBS (referred to as growth media). SK23MEL and HCC1954 cell lines were cultured in antibiotics-free RPMI 1640 (Wisent) supplemented with 10% FBS. SKOV3 cells were cultured in OSE (Wisent) supplemented with 10% FBS and 1% pen/strep antibiotics. Mouse embryonic fibroblasts (MEF; generated and immortalized as described in method details) were cultured in DMEM (Wisent) supplemented with 10% FBS and 1% pen/strep antibiotics. UWB1.289+BRCA1 cells were culture in 50/50 RPMI (Wisent) and MEGM (Lonza) media supplemented with 3% FBS and 200μg/mL G-418. All cell lines were negative for mycoplasma (Lonza, MycoAlert). Pyruvate-, glucose- and glutamine-free DMEM media (Wisent) is defined as assay media and supplementations of the media are described in method details depending on the experiment.

### Animal experimentation

NOD-*Rag1^null^ IL2rg^null^* (NRG; Jackson Laboratory) and CD73^-/-^ C57BL/6J mice (obtained from Dr. Linda Thompson; OMRF, Oklahoma City, OK, USA) were bred and housed at the CRCHUM which holds a certificate of Good Animal Practice from the Canadian Council on Animal Care (CCAC). C57BL/6J mice were purchased from Jackson Laboratory and housed at the CRCHUM. All the experimental procedures were authorized by an Institutional Animal Care and Use Committee (CIPA). For in vivo tumor growth, 8-12 weeks-old healthy female NRG mice were injected with 2×10^6^ MDA-MB-231 cells sub-cutaneously and tumor growth was monitored thrice weekly. Tumor volume was measured by caliper in two dimensions, and volumes were estimated using the equation V = (large diameter × small diameter^2^) × 0.5. The total number of mice analyzed for each experiment is detailed in figure legends. C57BL/6J and CD73^-/-^ C57BL/6J mice were used for mouse embryoblastic fibroblast (MEF) derivation (see cell lines generation and culture section).

### Flow cytometry

Single cell suspensions were strained (40μM nylon mesh) and incubated with fluorescence-conjugated antibodies (1:200 PE-Cy7 rat anti-mouse CD73 or 1:200 BV421 mouse anti-human CD73) for 30 minutes at 4°C in ice-cold PBS + 2% FBS + 2 mM EDTA (FACS buffer). Fluorescence was acquired with a LSRFortessa flow cytometer (BD) or sorted with a FACSAria III cell sorting system (BD) equipped with FACSDiva software (BD). Dead cells were excluded using a fixable viability dye eF506 (Thermo Fisher Scientific; cat.: 65-0866-14). FACS data were further analyzed with FlowJo software (version 10.8.0).

### Glucose incorporation assay

Cells were cultured in assay media supplemented with 10% FBS and 2 mM glutamine (Sigma) for 30 minutes before being harvested, washed with PBS 1X twice and incubated with 100 μM of 2-NBDG (Cayman Chemical) for 30 minutes at 37°C in a 5% CO2 humid atmosphere. Cells were washed twice with PBS 1X and resuspended in FACS buffer (see flow cytometry section). Fluorescence was analyzed in the FITC channel on a LSRFortessa flow cytometer (BD).

### ALDH assay

Tumor-initiating cells were sorted based on ALDH activity and tested on seahorse analyzer as previously described (45) using the ALDEFLUOR^™^ kit (Stemcell). Briefly, cells were harvested and resuspended at 2.5×10^6^ cells/mL of ALDEFLUOR^™^ assay buffer. 5 μL/mL of the ADLEFLUOR^™^ reagent was added to each cell line test tubes and 0.5 mL per test tubes were immediately aliquoted in control tubes in which 10μL of ALDEFLUOR^™^ DEAB reagent were added. Cells were incubated for 30 minutes at 37°C washed and resuspended in ALDEFLUOR^™^ assay buffer and rapidly sorted on a FACSAria III cell sorting system (BD). 5×10^4^ freshly sorted ALDH^hi^ cells were seeded in seahorse XF24 plate in growth media and incubated overnight before seahorse analysis as described in analysis of extracellular flux section.

### Proliferation assays

2.5×10^3^ cells were seeded in a 48-well plate and cultured in growth media for 8h before being washed and switched to assay media supplemented with 10% FBS and 0.1-2 mM glutamine (Glutamax; Gibco) and/or 2.5-25 mM glucose (Sigma). The moment at which the media was changed is defined as baseline (day 0). Percentage of well confluency was measured by applying a phase mask over 4X photos (2 photos per well) taken every 2h for 5 days using the Incucyte technology. For each independent experiments, experimental conditions were tested in triplicate. Proliferation was calculated by reporting the confluence level every 0.5 days relative to confluence level at baseline for each well.

### Stable isotope tracing analysis (SITA)

For SITA, 3×10^6^ cells/well were seeded in a 6-well plate and cultured in 2 mL assay media supplemented with 10% dialyzed FBS (Wisent) and either 25 mM U-[^13^C]-glucose (Cambridge Isotope Laboratories) and 4 mM glutamine (Sigma) or 4 mM U-[^13^C]-glutamine (Cambridge Isotope Laboratories) and 25 mM glucose (Sigma) for 3h at 37°C. Metabolites were extracted from cells using dry ice-cold 80% methanol, followed by sonication and removal of cellular debris by centrifugation at 4°C. Metabolite extracts were dried, derivatized as tert-butyldimethylsilyl esters, and analyzed via GC-MS by the metabolomic core from the *Goodman cancer research centre* (McGill University). Labeled metabolite abundance was expressed relative to the internal standard D27 (D-myristic acid) and normalized to protein content. Mass isotopomer distribution was determined using a custom algorithm developed at McGill University (46).

### LC-MS analyses

For intracellular measurements of AMP, ADP, ATP, NAD+ and aspartate, 3×10^6^ cells were seeded in 100 mm petri dishes. Cells were incubated with drugs (7G2 mAb, rhADA or NR) for 48 hours in assay media supplemented with 25 mM glucose, 2 mM Glutamax and 10% dialyzed FBS. Media was washed once with PBS 1X, remove completely and cell culture dishes were quickly snap frozen on liquid nitrogen. Cells were scraped on ice and collected in 675 μL ice-cold extraction buffer [80% (vol/vol) methanol, 2 mM ammonium acetate, pH 9.0, with 10 μM AMP-13C10,15N5 (IS-AMP) as internal standard], transferred into polypropylene (PP) tubes, and sonicated in a cup-horn sonicator at 150 W for 2 min (cycles of 10s on, 10s off) in an ethanol/ice bath. Cell extracts were centrifuged at 4°C for 10 min at 25,830 × *g*, and supernatants were collected in ice-cold 2-mL PP tubes, to which 250 μL water was added. Polar metabolites were extracted with 1,080 μL of chloroform:heptane (3:1 vol/vol) by 2 × 10s vortex followed by 10-min incubation on ice and 15-min centrifugation at 4°C, 12,500 × *g*. From the upper phase, 600 μL was collected without carrying out any interface material and transferred into new cold 2-mL PP tubes. These tubes were centrifuged again, and 400 μL supernatant were collected into cold 1.5-mL PP tubes. Samples were frozen in liquid nitrogen and dried in two steps – first, in a SpeedVac Concentrator for ~2 h (maximal vacuum, no heat; Savant) at 4°C to remove methanol; and second, by lyophilization for 90 min (Labconco FreeZone) – and then stored at −80°C until used. Samples were reconstituted in 14 μL of Milli-Q water, and injections of 3 μL were performed in duplicate on an electrospray ionization LC-MS/MS system composed of an Agilent 1200 SL device (for LC) and a triple-quadrupole mass spectrometer (4000Q TRAP MS/MS; Sciex). Samples were separated by gradient elution of 12 min on a Poroshell 120 EC-C18, 2.1 × 75-mm, 2.7-μm column (Agilent Technologies) using mobile phase consisting of an aqueous solvent A (10 mM tributylamine, 15 mM acetic acid, pH 5.2) and an organic solvent B (95% (vol/vol) acetonitrile in water, 0.1% formic acid), at a flow rate of 0.75 mL/min and column oven temperature of 40°C. Quantification was performed as described in (47).

For extracellular measurements of NAD+, NMN and NR, 0.2 μg/mL of human recombinant CD73 or 5×10^4^ cells seeded in a 96-wells plate were cultured in assay buffer (2 mM MgCl_2_, 125 mM NaCl, 1 mM KCl, 10 mM Glucose, 10 mM HEPES pH 7.2, ddH_2_O) with or without exogenous NMN (0.2 mM) for 1-8hrs. Supernatant was collected in 40% methanol+40% acetonitrile dry ice-cold extraction solution (2:2:1 methanol:acetonitrile:aqueous ratio). Samples were flash frozen on dry ice and stored at −80°C. For LC-MS/MS analysis, samples (500μL) were thawed and AMP-13C10,15N5 (IS-AMP) was added as internal standard. Samples were incubated 1h at 4°C with vigorous agitation and centrifuged 10 min, 20K × *g*, 4°C. Supernatants were transferred to new polypropylene tubes, dried down at 10°C by centrifugal vacuum evaporation, then reconstituted in 50μL 80% acetonitrile in water before LC-MS/MS analysis. Samples (5μL injections) were separated by HILIC (Nexera X2, Shimadzu) at 0.25 mL/min using a gradient elution (A=MilliQ water, B=90% acetonitrile in water, both A and B containing 10mM ammonium acetate pH 9.0 and 5μM Agilent’s deactivator additive; gradient: 0 min 90% B, 2 min 90% B, 12 min 60% B, 15 min 60% B, 16 min 90% B (re-equilibration), 24 min 90% B) on a Poroshell 120 HILIC-Z 2.1 × 100 mm, 2.7 μm HPLC column (Agilent) following a guard column of similar material, both kept at 30°C. Metabolites were detected after ESI on a triple quadrupole mass spectrometer (QTRAP 6500, SCIEX) with polarity switching looking for the following transitions: negative ions IS-AMP 360.8/79.0, NAD 661.8/540.0; positive ions NR 255.0/123.1, NMN 334.9/123.2. Depending on the experiment, results normalized on protein content are shown as area of peak obtain by mass spectrometry analysis or absolute concentrations. For quantification of absolute concentration, a mix of pure standards prepared in 80% acetonitrile in water was used to build a calibration curve. No traces of NAD, NMN or NR were found in the assay buffer.

### Extracellular flux analysis

Oxygen consumption rate (OCR) and extracellular acidification rate (ECAR) were measured in real-time using Seahorse (XF24 and XFe96) extracellular flux analyzers (Agilent). Cells were seeded in XF24 (5×10^4^) or XFe96 (1×10^4^) and incubated between 12- and 48-hours depending on the experiment. For assays with 7G2 mAb, SCH58261, PSB1115, rhADA, NECA or ADO, drugs were added to the media after seeding and changed after 24h with fresh media. The day of the assay, cells were washed and incubated for 60 min in a CO_2_-free incubator in seahorse assay base media (Agilent) containing 25mM glucose (Sigma) and 2 mM glutamine (Sigma). For mitochondrial stress test, ATP synthase was inhibited by injection of 1 μM oligomycin (Sigma), followed by 1 μM FCCP (Sigma) to induce mitochondrial uncoupling to determine the spare/maximal respiratory capacity. Non-mitochondrial respiration was determined after rotenone (Sigma; 1 μM)/antimycin A (Sigma; 0.1 μM) injection. For glycolysis measurements, cells were glucose-starved for 60 minutes in seahorse assay base media containing 2 mM glutamine (Sigma). Glycolysis was measured upon glucose injection (Sigma; 25 mM) and glycolytic reserve upon oligomycin (Sigma; 1 μM) injection. Upon completion of the assay, data were normalized by cell counts (XF24) or crystal violet (XFe96) to correct for seeding density and analysis was performed using Seahorse Wave software (2 min mix, 2 min wait, 2 min measurements).

### Genome-wide transcriptional analysis

Total RNA was isolated from MDA-MB-231 cells using the RNeasy Plus Mini Kit (Qiagen) according to the manufacturer’s instructions. RNA was quantified using a DS-11 spectrophotometer (DeNovix). Samples were sent to *Génome Québec* (Montréal, QC, CA) for gene expression analysis using an Affymetrix microarray chip. Expression values were computed using the robust multi-array analysis (RMA) normalization method (‘affy’ package in Bioconductor) (48, 49). When multiple probe sets mapped to the same official gene symbol, we computed their average value. Differential expression (DE) analysis was performed using ‘limma’ (50) standard analysis pipeline. Significant DE gene were defined as genes with absolute fold-change ≥ 1.5 and adjusted P-value ≤ 0.05. Statistical analyses were performed between *NT5E* mRNA expression and genes expression using Spearman correlation.

### Quantitative real-time PCR analysis

Total RNA was isolated and quantified as described in genome-wide transcriptional analysis section. Reverse transcription of 1μg RNA was performed using the SuperScript^™^ VILO^™^ cDNA Synthesis Kit (Thermo Fisher Scientific) and CD73 or NRK1 gene expression was analyzed using Taqman probes for mouse *Nt5e* (Thermo Fisher Scientific; Mm00501910_m1) or human *NMRK1* (Thermo Fisher Scientific; Hs00944470_m1) relative to mouse *Actb* (Thermo Fisher Scientific; Mm00607939_s1) or human *ACTB* (Thermo Fisher Scientific; Hs99999903_m1) using the StepOne PCR machine (Thermo Fisher Scientific) and software (version 2.3).

### Western Blot

Adherent cells were washed with ice-cold PBS and lysed and scrapped in CelLytic^™^ M buffer (Sigma) with 1X Halt^™^ protease and phosphatase cocktail inhibitors (Thermo Fisher Scientific) before being centrifuged for 15 min at 20,000g at 4°C. Proteins were harvested from supernatant and quantified by using Bradford protein assay dye reagent (Bio-Rad). 25 μg of protein from whole cell extract were loaded in 4-10% acrylamide gels and transferred on nitrocellulose membranes. Membranes were stained overnight in 5% BSA-containing PBS Tween 0.1% with following antibodies: mouse anti-β-actin (1:5000), mouse anti-PLCγ (1:2000), mouse anti-CD73 (human; 1:1000), mouse anti-FLAG (1:1000) and rabbit anti-Cas9 (S. pyogenes; 1:1000). For PARylation assay, 1×10^6^ cells were seeded in 6-well plates and drugs was added in media for 48 hours. Before proceeding to protein extraction, cells were treated with 2 mM H_2_O_2_ for 20 minutes at room temperature. 50μg proteins were load in a 4-10% acrylamide gels, transferred on nitrocellulose membrane and stained overnight in 5% milk-containing PBS Tween 0.1% with mouse anti-PAR (1:1000). Proteins were revealed with fluorescent secondary anti-rabbit or anti-mouse antibodies (1:10 000; LI-COR) using the LI-COR fluorescent scanner or by chemiluminescence (Bio-Rad) using HRP-conjugated secondary anti-mouse antibody (2.5:10 000) and Amersham ECL prime detection reagent (GE healthcare lifescience).

### Immunostaining

For γH2AX immunofluorescence staining, 5×10^4^ cells were seeded on glass slides (Thermo Fisher Scientific) and incubated overnight before adding treatments (Doxorubicin, carboplatin, olaparib). 48h later, cells were fixed with formalin 10% (Sigma) for 10 min at room temperature, washed with PBS 1X and then permeabilized with 0.25% Triton-X100 (Sigma) in PBS 1X for 15 min. Slides were incubated 1 hour with DAKO blocking solution (Agilent) and incubated overnight at 4 °C with anti-phospo-H2AX (1:2000). Cells were then washed 3 times with PBS 1X-Tween 0.05% followed by incubation with the secondary antibody (1:800; donkey anti-mouse IgG Alexa Fluor 488; Thermo Fisher Scientific). The glass slides were stained with DAPI, washed, and mounted with Fluoromount^™^ aqueous mounting medium (Sigma). Stained sections were scanned (Leica) by the molecular pathology core of the CRCHUM and photographs were analyzed using Visomorph^™^ viewer software for automatic counting of γH2AX foci.

### Data representation and statistical analysis

Mean +/- SEM are shown. Each experimental conditions for all independent experiments were tested in duplicates or triplicates excepted for XFe96 extracellular flux analysis (n=4-6 per condition tested). The numbers of repeated independent experiments are indicated in figure legends. Student T-test were performed when comparing 2 conditions. 1-way ANOVA was used when comparing 3 or more conditions. Statistical significance is shown in comparison to the control condition (untreated or CD73+), unless indicated otherwise by a bracket. Results are considered significant when *P* < 0.05. All statistical analyses were performed using graph pad prism software (version 9.0.2).

## Material and data availability

Further information and requests for resources, reagents and cell lines (supplementary table 1) generated in this study should be directed to and will be fulfilled by the lead contact, John Stagg. This study did not generate new unique reagents. Any additional information required to reanalyze the data reported in this study will be made available from the lead contact upon request.

---

**Table 1.**
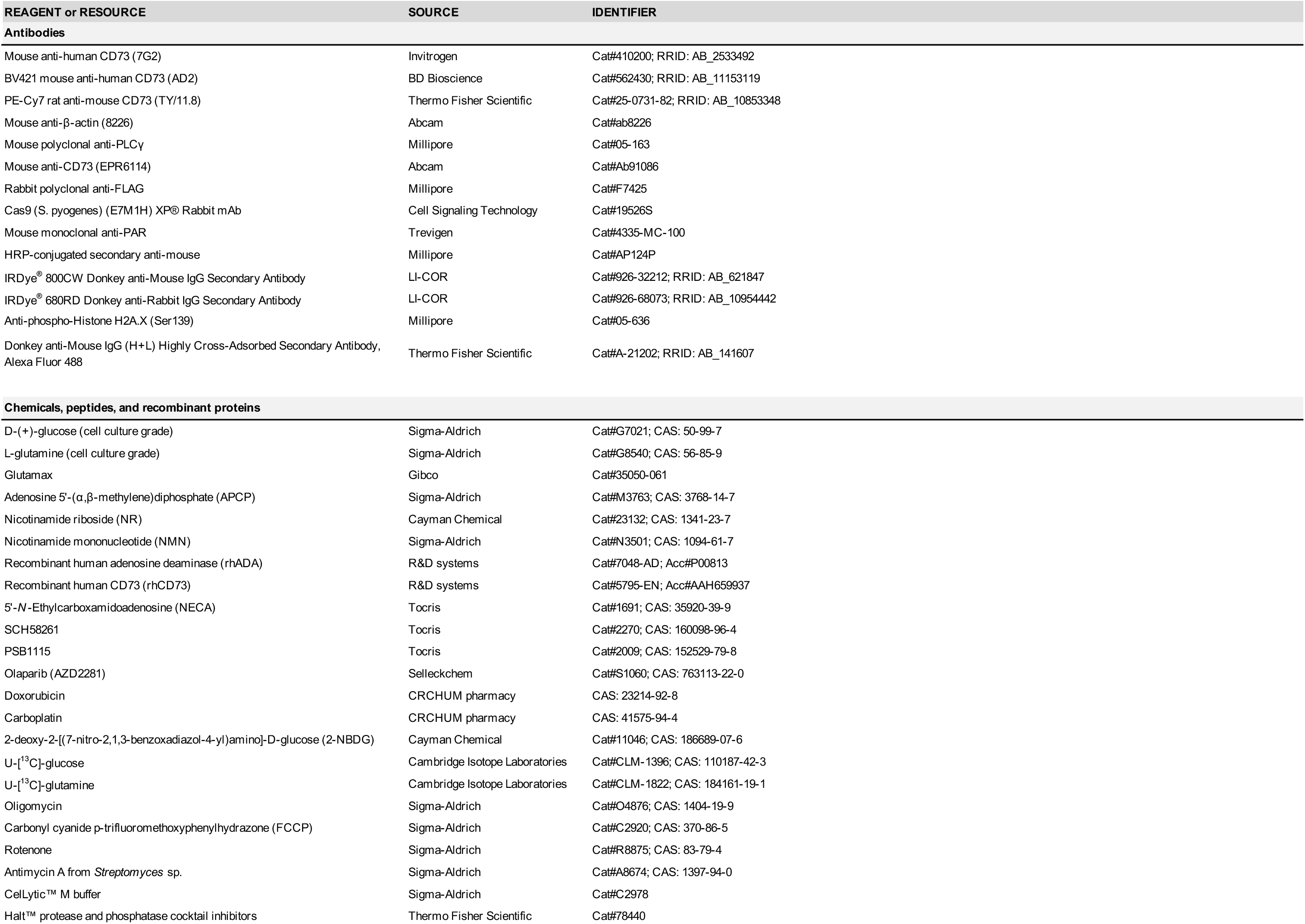
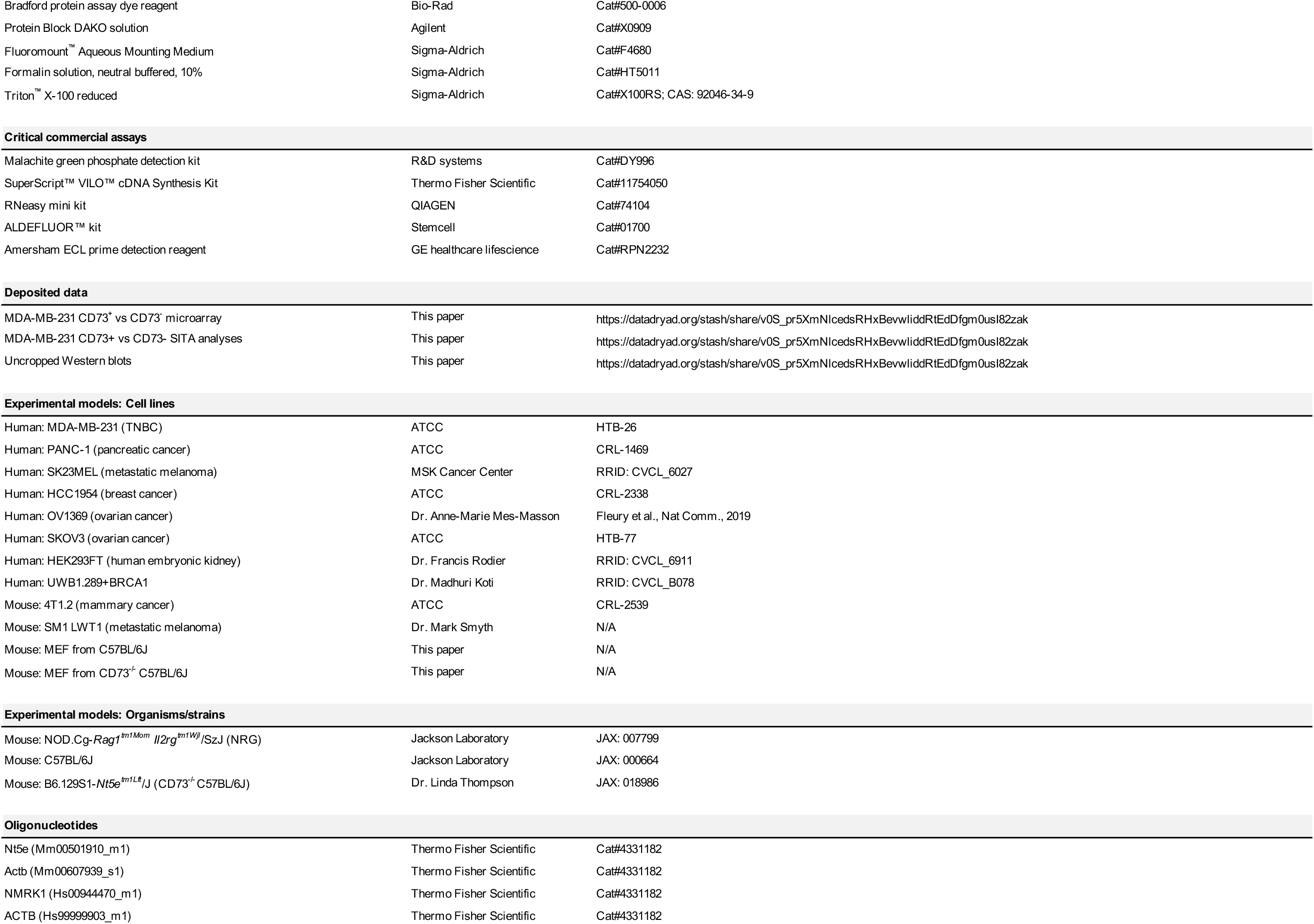
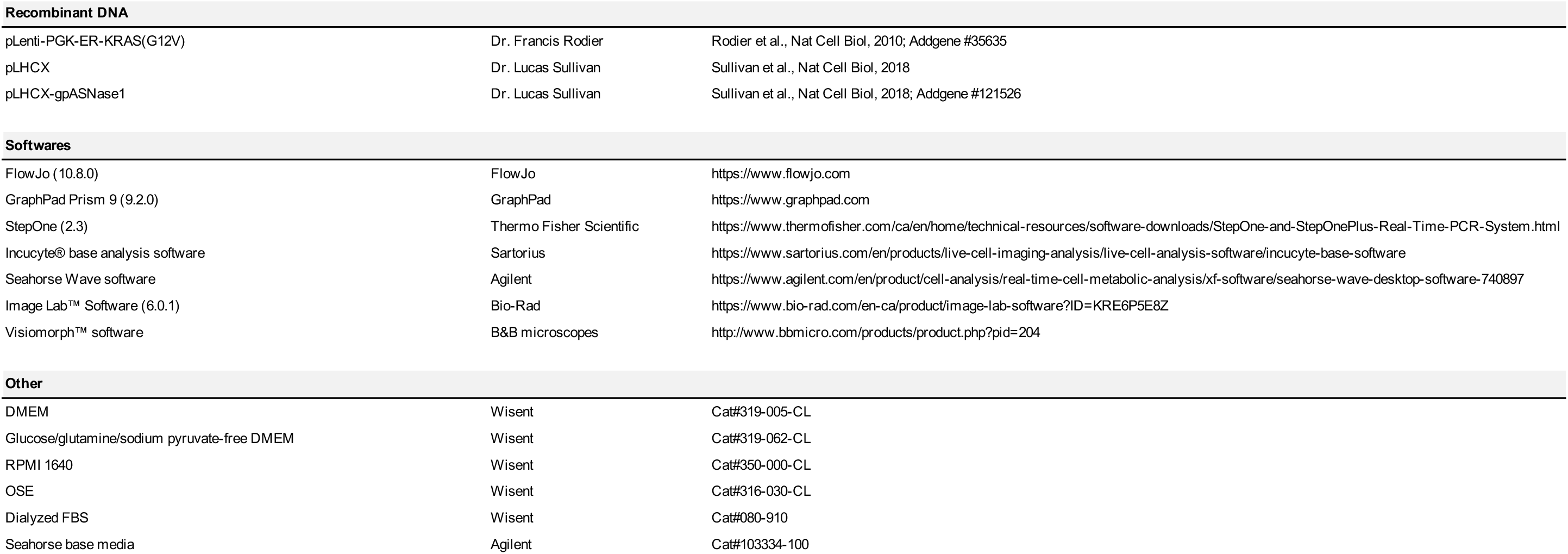
List of reagent and resources with unique identifier.

## SUPPLEMENTARY FIGURES

**Figure S1.**
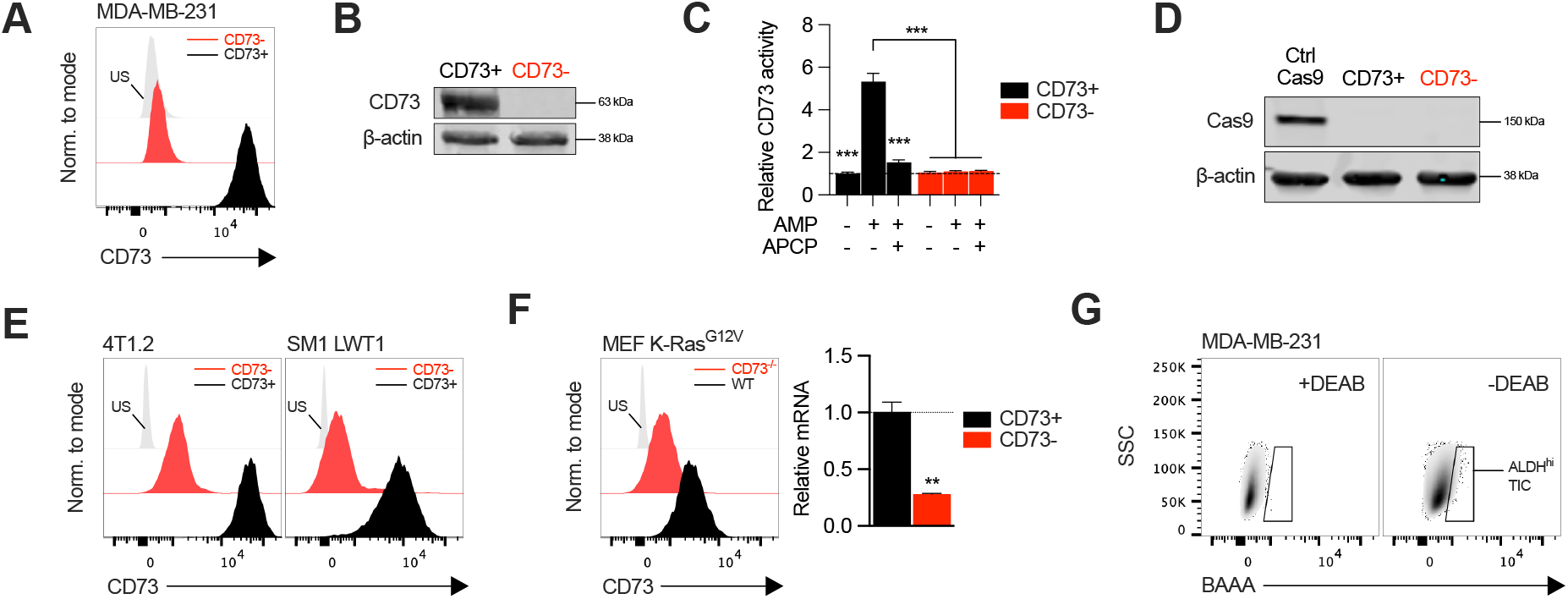
Generation of CD73-knockout cell lines. **A-C** Knockout efficiency of CD73 on MDA-MB-231 cells following CRISPR-Cas9 gene edition. Cell sorting enrichment was confirmed by cell surface (**A**; FACS. US=unstained control) and total (**B**; western blot) CD73 expression. **C** Analysis of CD73 hydrolyzing enzymatic activity using malachite green assay compared between CD73+ and CD73-MDA-MB-231 cultures. APCP=CD73 inhibitor. **D** Western blot validating transient Cas9 expression in MDA-MB-231 cells. **E** Knockout efficiency of CD73 on 2 mouse cancer cell lines (4T1.2 and SM1 LWT1) following CRISPR-Cas9 gene edition. Cell sorting enrichment was confirmed by FACS. **F** Mouse embryoblastic fibroblasts (MEF) were derived from WT and CD73^-/-^ C57BL/6 mice and transformed with an oncogenic K-Ras in vitro. Emerging transformed clones were subsequently isolated by sub-culture and analyzed for CD73 surface protein expression and *Nt5e* gene expression (n=2). **G** Gating strategy for MDA-MB-231 TIC sorting based on ALDH-mediated conversion of fluorescent BAAA. The gating is defined in presence of DEAB according to manufacturer protocol, an inhibitor of ALDH enzymatic activity. Means +/- SEM are shown (**p<0.01; ***p<0.001 by Student T tests).

**Figure S2.**
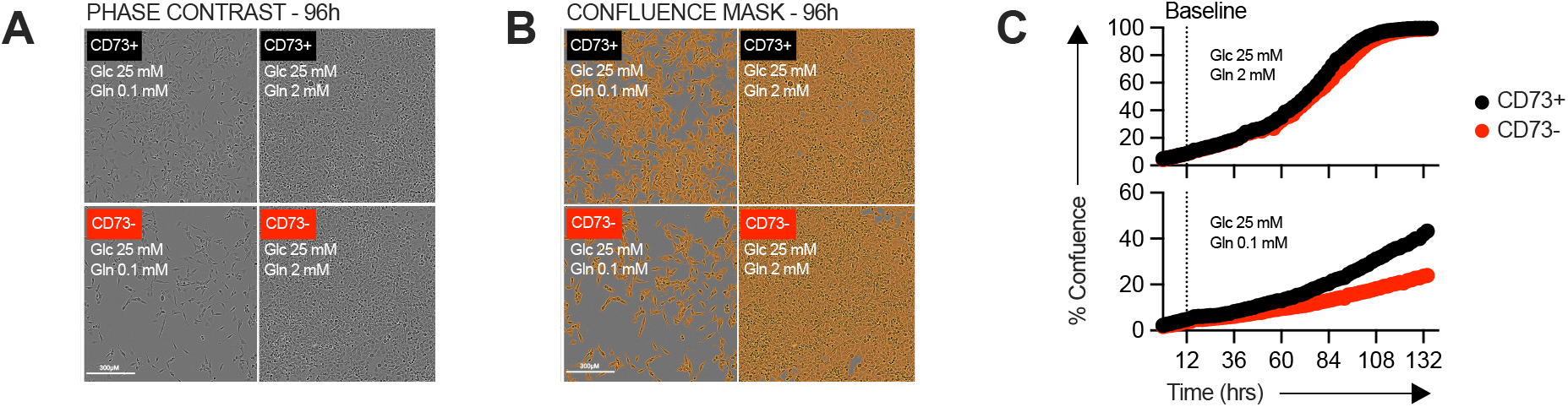
Proliferation assay using Incucyte live imaging technology. **A-B** Examples of in vitro proliferation imaging in presence of excess (2 mM) or limited (0.1 mM) amount of glutamine (Gln) measured using Incucyte technology. Proliferation was quantified by measuring confluence level at different timepoints. **A** Phase contrast representative images of CD73+ and CD73-MDA-MB-231 cell cultures 72h after initiation of assay. **B** Same images (as in **A**) with a confluence mask applied. **C** Percentage of well confluence over time (0-5 days). Cells were seeded in regular complete media for 8-12h before being switched to assay media (excess or limited Gln), which is defined as baseline confluence (dashed line). Proliferation index (**Fig. 1B**) is expressed as a ratio of increased confluence every 0.5 days relative to baseline.

**Figure S3.**
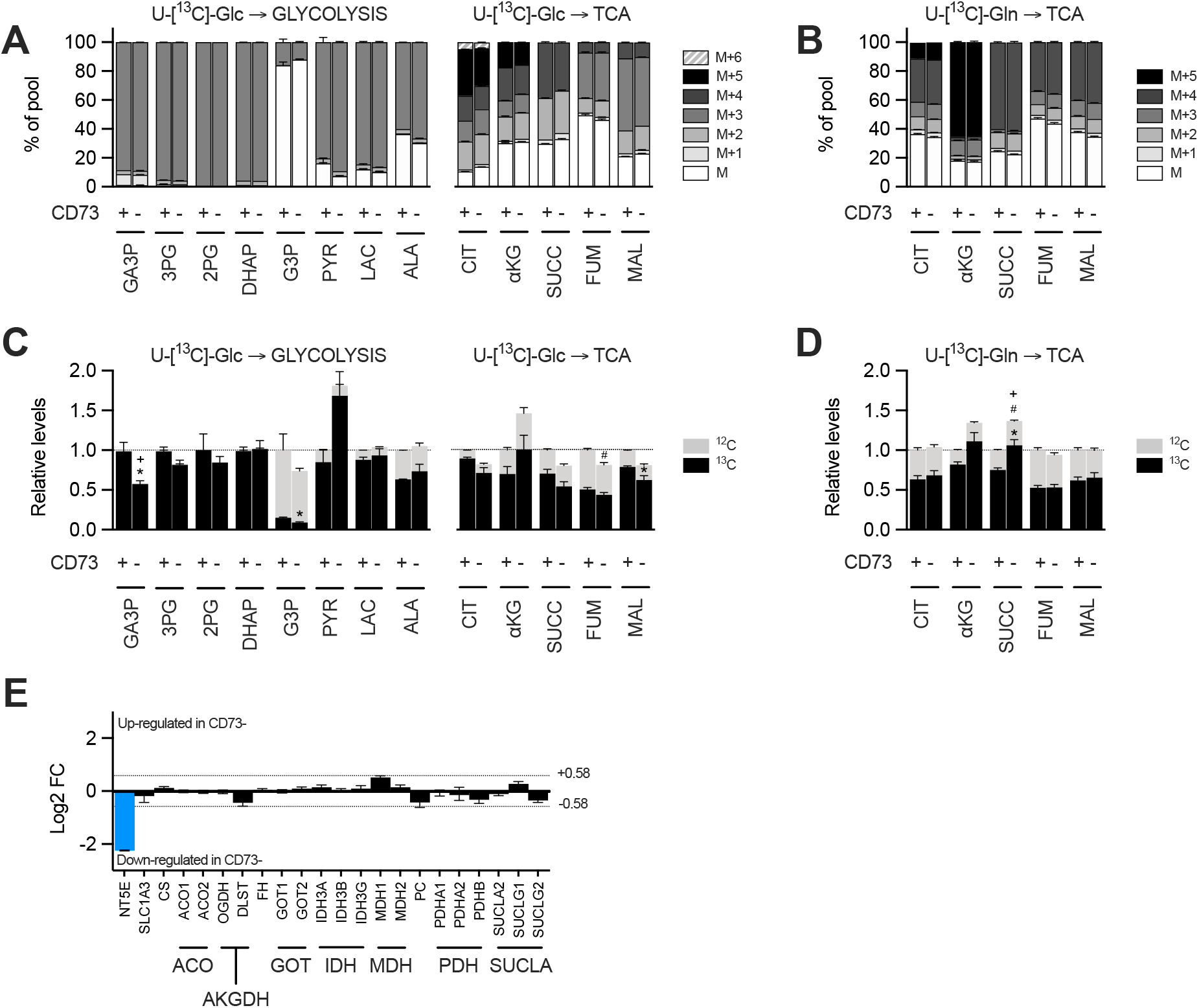
Glucose and glutamine SITA and transcriptional analysis of TCA cycle enzymes in MDA-231 cells. **A-D** Stable isotope tracer analysis (SITA) of MDA-MB-231 cells cultured with U-[^13^C]-glucose or U-[^13^C]-glutamine for 3h in vitro. Isopotomer distribution of U-[^13^C]-glucose (**A**) and U-[^13^C]-glutamine (**B**) in metabolites from glycolytic and TCA cycle pathways compared between CD73+ and CD73-cells. Glucose (**C**) and glutamine (**D**) contribution to glycolytic and TCA intermediates. Metabolites levels (**C-D**) were normalized on cell number and are shown relative to CD73+ cells (n=1 per tracer). Stats: *comparison of ^13^C levels; ^#^Comparison of ^12^C levels; ^+^Comparison of total ^13^C+^12^C levels. **E** Transcriptional analysis by microarray of TCA enzymes showing gene regulation (Log2 of fold change; FC) in CD73-compared to CD73+ MDA-MB-231 cells. Dashed line show threshold of 1.5-fold change (+/- 0.58 Log2 of FC). CD73 (NT5E; blue bar) is shown as an internal control (n=1). Means +/- SEM are shown (*p<0.05 by Student T tests).

**Figure S4.**
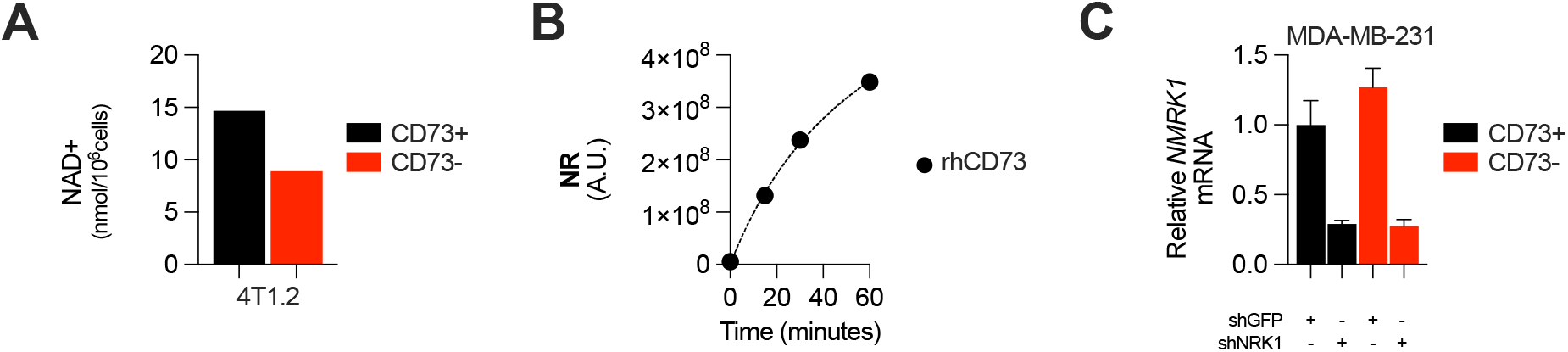
Complement to figure 3: CD73 contributes to metabolic fitness of cancer cells through NAD synthesis. **A** LC-MS analysis of intracellular nicotinamide adenine levels (NAD+) normalized to protein content of CD73pos (CD73+) and CD73neg (CD73-) 4T1.2 cells (n=1). **B** LC-MS analysis of nicotinamide riboside (NR) production over 60 minutes in supernatant of recombinant protein hCD73 incubated with 0.2 mM nicotinamide mononucleotide (NMN). NR levels are shown as area under peak by mass spectometry analysis (n=1). **C** NRK1 mRNA expression analyzed by qPCR in shGFP- and shNRK1-transfected MDA-MB-231 CD73+ and CD73-cells.

**Figure S5.**
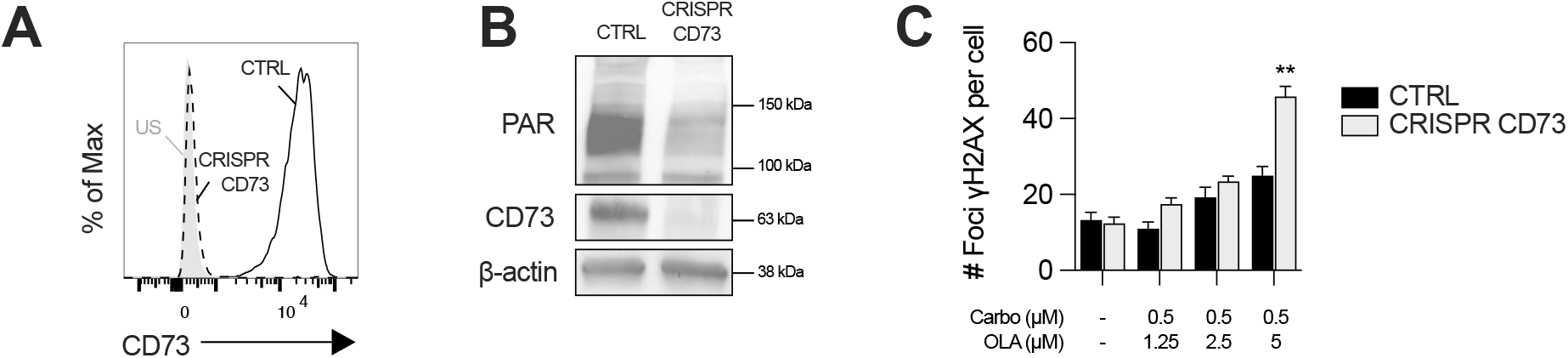
CD73 deficiency sensitizes UWB1.289+BRCA1 cells to DNA-damaging agents. **A** Knockout efficiency of CD73 expression on UWB1.289+BRCA1 cells analyzed by FACS. US=unstained control. **B** Western blot showing CD73 protein levels and PARylation levels in CD73+ (CTRL) and CD73- (CRISPR CD73) UWB1.289+BRCA1 cells (n=2). β-actin is used as a loading control. **C** Number of γH2AX foci per CD73+ (CTRL) and CD73- (CRISPR CD73) UWB1.289+BRCA1 cells treated with carboplatin (0.5μM) and olaparib (1.25-5μM) for 48h (n=2). Means +/- SEM are shown (**p<0.01; by Student T tests).

